# Combination of Haloperidol with UNC9994, β-arrestin-biased analog of Aripiprazole, ameliorates schizophrenia-related phenotypes induced by NMDAR deficit in mice

**DOI:** 10.1101/2024.04.25.591166

**Authors:** Tatiana V. Lipina, William C Wetsel, Marc G. Caron, Ali Salahpour, Amy J. Ramsey

## Abstract

**Background:** Glutamatergic system dysfunction, particularly involving the N-methyl-D-aspartate receptor (NMDAR), contributes to a full spectrum of schizophrenia-like symptoms, including the cognitive and negative symptoms that are resistant to treatment with antipsychotic drugs (APDs). Aripiprazole, an atypical antipsychotic drug (APD), acts as a dopamine partial agonist and its combination with haloperidol (a typical APD) has been suggested as a potential strategy to improve schizophrenia symptoms. Recently, an analog of aripiprazole - UNC9994 was developed. UNC9994 does not affect D2R-mediated Gi/o protein signaling but acts as a partial agonist for D2R/β-arrestin interactions. Hence, our objective was to probe the effects of co-administrating haloperidol with UNC9994 in NMDAR mouse models of schizophrenia.

**Methods:** NMDAR hypofunction was induced pharmacologically by acute injection of MK-801 (NMDAR pore blocker; 0.15 mg/kg) and genetically by knockdown of Grin1 gene expression in mice, which have a 90% reduction in NMDAR levels (Grin1-KD). After intraperitoneal injections of vehicle, haloperidol (0.15 mg/kg), UNC9994 (0.25 mg/kg) or their combination mice were tested in open field, Pre-Pulse inhibition (PPI), Y-maze and Puzzle box.

**Results:** Our findings indicate that low dose co-administration of UNC9994 and haloperidol reduces hyperactivity in MK-801-treated animals and in Grin1-KD mice. Furthermore, this dual administration effectively reverses PPI deficits, repetitive/rigid behavior in the Y-maze, and deficient executive function in the Puzzle box in both animal models.

**Conclusions:** The dual administration of haloperidol with UNC9994 at low doses represents a promising approach to ameliorate positive, negative, and cognitive symptoms of schizophrenia.

**Significance statement:** Schizophrenia is a devastating mental disorder and characterized by positive, negative, and cognitive symptoms. Cognitive and negative symptoms remain a focus of research dedicated to development of effective antipsychotic drugs (APDs). Aripiprazole, an atypical APD, acts as a dopamine partial agonist and its combination with haloperidol (a typical APD) has been suggested as a potential strategy to improve schizophrenia symptoms. An analog of aripiprazole - UNC9994 was recently developed, which does not affect D2R-mediated Gi/o protein signaling but acts as a partial agonist for D2R/β-arrestin interactions. Our pre-clinical findings on pharmacological (MK-801, 0.15 mg/kg) and genetic (Grin1-KD) mouse models of NMDAR deficiency showed that the dual administration of UNC9994 (0.25 mg/kg) with haloperidol (0.15 mg/kg) at low doses reduces hyperactivity, corrects prepulse inhibition (PPI) deficits, rigid behavior in the Y-maze, and deficient executive function in the Puzzle box. Further studies of the polypharmacy of UNC9994 with APDs is essential to facilitate translational studies in clinics.

## Introduction

Schizophrenia is a devastating mental disorder affecting nearly 1 % of the population and accounting for 25 % of psychiatric hospital beds. Its symptomatic structure includes positive, negative, disorganized, excited, and depressed factors (van der Gaag et al., 2006; Wallwork et al., 2012). Cognitive impairments are a core of schizophrenia (Elvevåg et al., 2000) as they are observed among all subtypes of schizophrenia (Heinrichs et al., 1993). The disorganized factor (deficient abstract thinking, attention, working memory, learning, delayed processing speed, disorientation, stereotyped thinking, and conceptual disorganization) is strongly linked to cognitive test scores (Rodriguez-Jimenez et a., 2013).

Cognitive, and negative symptoms respond minimally to currently available antipsychotic drug treatments (APDs) (Miyamoto et al., 2012), and therefore, these symptoms remain a focus of intense research dedicated to APDs development. In the 1990s, atypical APDs were introduced in the hope that they could ameliorate cognitive symptoms in addition to psychotic symptoms without side effects. However, it has been shown that all APDs elicit comparably modest effects on cognitive functions (Keefe, 2007). Besides APDs, other drugs have been re-purposed in attempts to correct some symptoms of schizophrenia, including e.g. anticholinergic drugs, antidepressants, treatments to manage obesity, type II diabetes, cardiovascular diseases, or sleep disturbances (McCutcheon et al., 2023), but their effectiveness is still unclear. Several conceptually novel treatments were introduced to alleviate some cognitive symptoms of schizophrenia, including e.g. Trace amine-associate receptor 1 (TAAR1) agonists (Koblan et al. 2020), M1/M4 muscarinic receptor agonist KarXT (Kaul et al., 2024), cannabidiol (McGuire et al., 2018), riluzole and memantine (Farokhnia et al. 2014), glycine transporter inhibitor (Fleischhacker et al., 2021) or luvadaxistat, a d-amino acid oxidase inhibitor (ACNP 60^th^ Annual Meeting, 2021). Although no pharmacological treatments have received official approval for preventing or treating the cognitive symptoms of schizophrenia, significant advancements in biomedical research may offer hope in addressing these challenges (Lobo et al. 2022).

The difficulty in designing APDs arises from our incomplete understanding of the molecular causative mechanisms of schizophrenia. Genetic studies point towards high polygenicity of schizophrenia suggesting that higher clinical efficacy might be achieved with multitarget drugs rather than single-target compounds (Hopkins, 2008). For instance, our previous study showed synergistic effect between phosphodiesterase 4B (PDE4) and glycogen synthase kinase-3 (GSK-3) inhibitors to correct schizophrenia-related behavioural endophenotypes in DISC1-L100P mutant mice (Lipina et al., 2011), supporting the idea of multitarget drugs in mental health disorders (Talevi et al., 2012).

All current APDs act via antagonism of the dopamine D2R [Miyamoto et al., 2012; Seeman and Lee, 1975; Kaar et al., 2020). In addition to the cAMP-dependent action, D2R activation also can affect other signalling pathways (Beaulieu et al., 2008), eliciting its effects in a G-protein-dependent and independent manner (Masri et al., 2008). G-protein-dependent action is a classical feature of GPCRs, but the G-protein-independent mechanism of D2R is mediated via b-arrestin (Beaulieu et al., 2005). Pharmacological activation of D2R in mice inhibits Akt kinase with concomitant activation of GSK-3 (Beaulieu et al., 2004). Both typical and atypical APDs enhance the phosphorylation of Akt and GSK-3, inhibiting GSK-3 enzymatic activity (Emamian et al., 2004). Thus, it was hypothesized that targeting β-arrestin signalling might be a promising direction for the development of new APDs (Allen et al., 2011). Indeed, a recent study generated three new unique compounds, UNC9975, UNC0006, and UNC9994 - D2R agonists that display a bias of β-arrestin signalling over Gi-coupled pathways signalling (Allen et al., 2011). Antipsychotic-like activity was reported for the UNC9994 compound in mice that was absent in β-arrestin-2 knockout mice. A follow-up pre-clinical study found robust antipsychotic-like effects of UNC9975 and UNC9994 on phencyclidine (PCP) pharmacological and Grin1 knockdown (Grin1-KD) genetic models of schizophrenia (Park et al., 2016).

All these unique compounds were generated through a robust diversity-oriented modification of aripiprazole which acts as a partial agonist of the D2 dopamine receptors like haloperidol. Its mixed agonist/antagonist action is thought to be responsible for its favourable efficacy. Aripiprazole also has fewer cardio-metabolic effects and hyperprolactinemia than other atypical APDs (Chrzanowski et al., 2006; Newcomer et al., 2008) and is superior in reducing positive, and possibly negative symptoms of schizophrenia with minimal risk of re-hospitalization (Pigott et al., 2003).

Antipsychotic *polypharmacy* (co-prescription of two or more APDs) is a common practice for treatment resistant schizophrenia. A recent study hypothesized that dual administration of aripiprazole with haloperidol could be beneficial for patients with schizophrenia (Crapanzano et al., 2022). The study reasoned that when aripiprazole is added to haloperidol, it should displace haloperidol from D2R binding sites and elicit its partial agonist intrinsic activity (Carboni et al., 2012), and therefore attenuate haloperidol-induced adverse effects and ameliorate negative symptoms (Shim et al., 2007; Mahgoub et al., 2015). However, it is possible the addition of haloperidol to aripiprazole may cause a powerful D2R inhibition and reduce aripiprazole’s beneficial effects. Nevertheless, a recent case report, indeed, demonstrated a beneficial effect of aripiprazole combined with haloperidol (Kuo and Hwu, 2008).

In this light, it could be suggested that the combination of haloperidol with the UNC9994 compound may also elicit specific and potent antipsychotic-like activity to correct schizophrenia-related cognitive and negative phenotypes. Therefore, our current study aimed to probe the effects of dual administration of haloperidol with UNC9994 on pre-clinical models of schizophrenia.

Animal models of schizophrenia are valuable tools for testing novel pharmacological strategies for the treatment of this disorder. MK-801-treated C57BL/6J mice, or Grin1-KD mice display hyperactivity, deficient working memory assessed in Y-maze, impaired filtration of irrelevant stimuli assessed by pre-pulse inhibition of acoustic startle response (PPI) as well as deficient executive function measured in the Puzzle box (Mohn et al., 1999; Milenkovic et al., 2014; Islam et al., 2017; Mielnik et al., 2021; Lipina et al., 2022).

Using these two pre-clinical models, we found that co-administration of haloperidol and UNC9994 in low doses reduced motor hyperactivity induced by MK-801 and agitation observed in Grin1 genetic knockdown in mice. Dual administration of haloperidol with UNC9994 also reversed PPI deficit, repetitive/rigid behaviour in Y-maze and improved executive function in the Puzzle box test. Our results show that dual administration of haloperidol with UNC9994 at low doses is an efficient approach to ameliorate schizophrenia-like phenotypes including positive, negative, and cognitive symptoms.

## 2. Materials and Methods

### 2.1. Animals

Knockdown mice for the NR1 subunit of the NMDA receptor (GluN1 knockdown; Grin1-KD) were generated in the animal facility of the University of Toronto as previously described (Mielnik et al., 2021). In Grin1-KD mice, there is an insertion of a neomycin cassette in the intron 10^th^ of the GRIN1 gene, flanked by loxP sites. Grin1^+^/^flneo^ C57Bl/6J congenics and Grin1^+^/^flneo^ 129×1/SvlmJ congenic mice were intercrossed to produce experimental mice [(Grin1+/+; wild type; WT); and Grin1KD)] mice as recommended (Silva et al., 1997) to minimize the confound of homozygous mutations on each parental strain. Grin1-KD mice express only 5-10% of normal levels of GluN1 subunit of the NMDAR complex. Two to five mice of mixed genotypes (WT; Grin1+/-; and Grin1-KD) were housed per cage with a 12 h light-dark cycle (7:00 a.m.–7:00 p.m.) with *ad libitum* access to food and water.

All experimental procedures were performed on naïve male and female WT and Grin1-KD mice between 12-16 weeks of age during the light cycle (9:00 a.m.–5:00 p.m.). MK-801 pharmacological studies were performed on adult WT mice of both sexes produced from the breeding of Grin1-KD mice.

Experimental mice were transported from the mouse colony room to the experimental room for 30-minutes of habituation before any behavioral procedure. All behavior was assessed by a skilled experimenter blind to genotype and drugs administration. All procedures were approved by the University of Toronto Faculty of Medicine and Pharmacy Animal Care Committee in compliance with the Animal Research Act of Ontario and the Guidelines of the Canadian Council on Animal Care.

### 2.2. Behavioral Tests

*Open Field:* Mice were placed in a Plexiglass chamber (20 × 20 × 45 cm3) for 60 minutes. Total distance travelled was measured in 5 min bins by infrared beam breaks using Versamax activity monitors (Omnitech Electronics, Columbus, OH, USA).

*Pre-pulse Inhibition (PPI) of Acoustic Startle Response (ASR):* Pre-pulse inhibition of the acoustic startle response was measured with SRLAB equipment and software from San Diego Instruments. Background white noise was maintained at 65 dB. PPI procedure had 80 randomized trials and was structured as follows: pulse alone (100 dB above background), pre-pulse alone (4, 8, or 16dB above background), pre-pulse plus pulse and no pulse. Five pulse-alone trials were performed at the start and end of the 80 trials. The onset of the pulse following the pre-pulse was delayed 100 ms. The time interval between trials was randomized from 5s-20s. Pre-pulse inhibition was measured as a decrease in the amplitude of startle response to a 110 dB acoustic startle pulse, following each pre-pulse (4, 8, and 16 dB). The percentage of PPI induced by each prepulse intensity was calculated as [1 – (startle amplitude on prepulse trial)/(startle amplitude on pulse alone)] * 100%.

*Y-maze:* The Y-maze task was performed as previously described (Lipina et al., 2013). The maze consists of a three-arms (labelled as arm “A”, arm “B”, and arm “C”) horizontal maze (40 cm x 8 cm x 15 cm) (Noldus Information Technology, The Netherlands) in which the arms are symmetrically disposed at 120° angles from each other. The maze floor and walls were constructed from white opaque polyvinyl plastic with distinctive geometric shapes on the walls. A mouse was initially placed in the start arm (arm “A”) and the sequence (i.e. ABCCAB etc) and number of arm entries were recorded for 5 min. Arm entries were defined as all four paws entering the arm. Spontaneous alternation refers to visiting all three arms in sequence (i.e. ABC or CAB but not CBC). The percentage of alterations was defined according to the following equation: % Alteration = [(Number of alterations)/(Total arm entries – 2)] * 100. The number of re-visits of the previously visited arm was manually scored as the index of repetitive/rigid behaviour. The total number of arm entries serves as an indicator of ambulation.

*Puzzle box:* The puzzle box test was performed as previously described with mild modifications (Lipina et al., 2022). Mice were tested on three trials over two consecutive days for a total of six trials (T1-T6). The puzzle box was a brightly lit arena (58 × 28 × 27.5 cm^3^) connected to a darkened goal box (14 × 28 × 27.5 cm^3^) by a wall divider. In trial T1 mice used a doorway and underpass to reach the goal box. In trials T2, T3, and T4 the doorway was blocked leaving an open underpass. In trials T5 and T6 the underpass was blocked with corncob bedding. This sequence of trials allowed assessing problem-solving ability (T2 and T5), and learning/short-term memory (T3 and T6), while the repetition after ∼ 24 h provided a measure of long-term memory (T4). Each trial started by placing the mouse in the start zone, and ended when the mouse entered the goal zone with all four paws, or after a total time of 3 minutes. The performance of mice in the puzzle box was assessed by measuring the latency to enter the goal zone.

### 2.3. Drugs

Haloperidol and MK-801 maleate salt were purchased from Sigma-Aldrich (St. Louis, MO). UNC9994 was provided by Dr. Jian Jin from the University of North Carolina in Chapel Hill. Haloperidol was dissolved in 0.3% Tween-20 (Bio-Rad) and brought to volume with saline 0.9% NaCl. UNC9994 was dissolved in a solution of 0.8% glacial acetic acid in 15% hydroxypropyl β-cyclodextrin in sterile water. MK-801 was dissolved in saline (0.9% NaCl). All compounds were administered intraperitoneally (i.p.) in a 5 ml/kg volume. WT mice were injected with MK-801 or vehicle and returned to their home cage for 10 minutes. Then, MK-801-treated or vehicle-treated WT mice were given either vehicle, haloperidol, UNC9994 alone or co-treated with haloperidol and UNC9994, and then either immediately placed for the behavioural testing in the open field, or after 20 minutes to test animals in PPI, Y-maze, or Puzzle box tests. As for Grin1-KD mice, drug-treated animals were immediately placed in the open field. UNC9994, haloperidol, and their combination were administrated to Grin1-KD mice with 30 minutes as a pre-treatment time to probe their effects in PPI, Y-maze, and Puzzle box. The doses of MK-801 (0.15 mg/kg), UNC9994 (0.25 mg/kg), and haloperidol (0.15 mg/kg) were determined from pilot experiments and based on other studies (Lipina et al., 2005; Duncan et al., 2006; Park et al., 2016).

### 2.4. Statistics

Statistical analyses were completed using TIBCO software (Statsoft, Dell) and Prism GraphPad (La Jolla, CA) software. Because there were no significant sex effects, data for both sexes were combined. Behavioural data were analyzed using two-way ANOVAs with repeated measures with the appropriate between-subjects and within-subjects factors and Pearson’s correlation analysis. Significant main effects or interactions were followed by Fisher’s least significant difference (LSD) post hoc comparisons.

## 3. Results

### 3.1. Effects of UNC9994 (0.25 mg/kg), haloperidol (0.15 mg/kg) and their co-administration on hyperactivity in MK-801 (0.15 mg/kg) treated and in Grin1-KD mice

#### 3.1.1. MK-801 pharmacological model

The total travelled distance assessed in the open field test, was affected by MK-801 treatment [F (1, 46) = 78.6; p < 0.0001], co-administered drugs (haloperidol, UNC9994 and haloperidol + UNC9994) [F (3, 46) = 6.8; p < 0.0001], and their interactions [F (3, 46) = 3.1; p < 0.05]. There was a significant effect of time intervals [F (11, 506) = 8.1; p < 0.001], and their interactions either with MK-801 [F (11, 506) = 21.3; p < 0.0001], drugs [F (33, 506) = 3.3; p < 0.001] or their interactions [F (33, 506) = 2.6; p < 0.001] on ambulation. As expected, MK-801 induced hyperactivity (p’s < 0.01-0.001) (Figure 1B). Administration of UNC9994 or haloperidol to MK-801-pre-treated mice modestly decreased their ambulation (p’s < 0.05-0.01) while the dual injection of haloperidol with UNC9994 compound reduced motor activity significantly (p’s < 0.01-0.001). At the doses used, haloperidol or UNC9994 administrated alone, or in combination did not affect vehicle-treated mice (Figure 1A, D).

**Figure 1.**
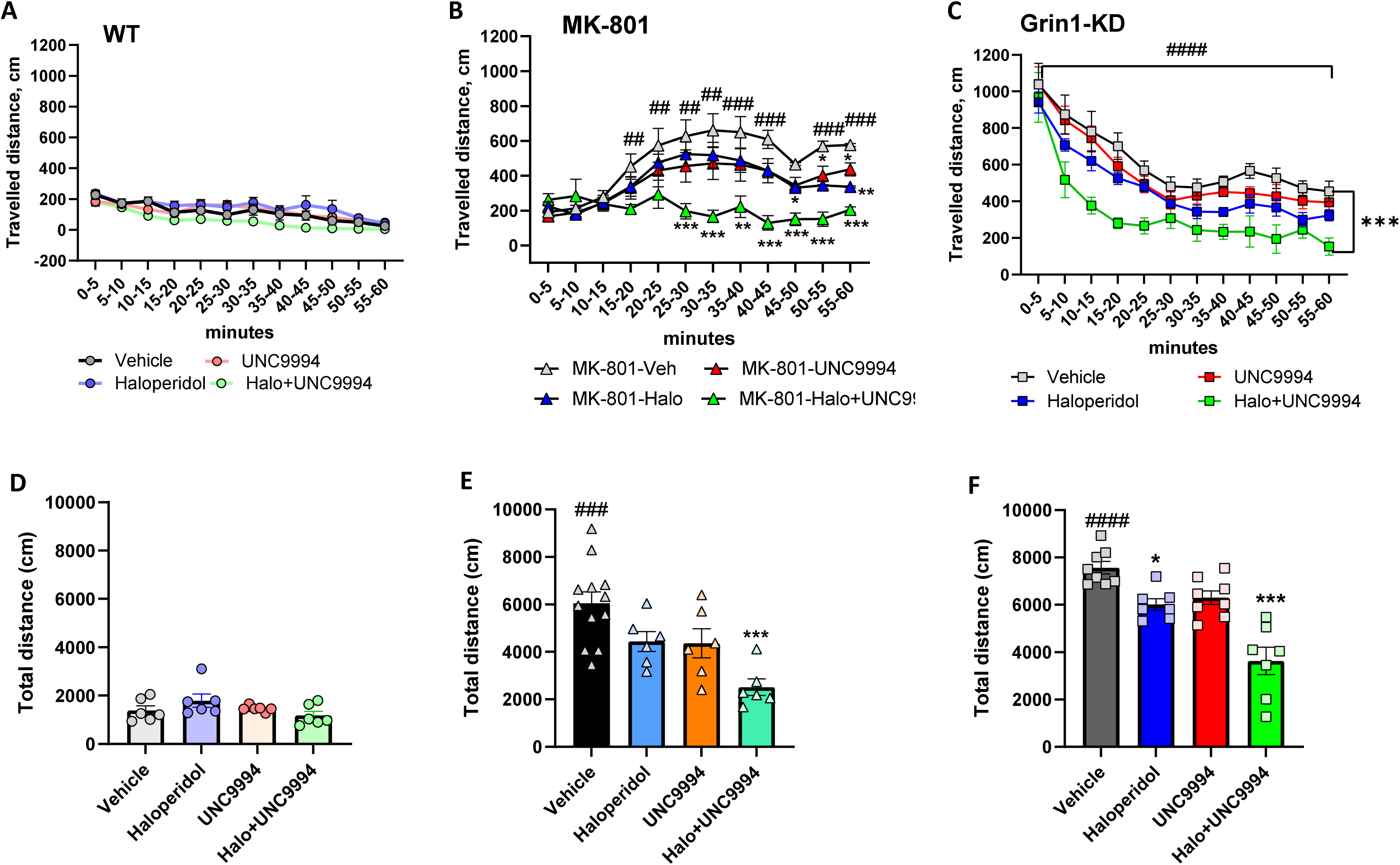
**A-F.** Effects of haloperidol (0.15 mg/kg), UNC9994 (0.25 mg/kg), and their co-administration on hyperactivity assessed in the open field for 60 minutes in vehicle-/WT (**A, D**), MK-801 (0.15 mg/kg)-treated wild-type (WT) mice (**B, E**), and Grin1 knockdown (Grin1-KD) mice (**C, F**). The travelled distance is presented with 5-minutes bin (**A-C**) and as total ambulation (**D-F**) for each experimental group: WT (n = 6/6/6/6); MK-801-treated mice (n = 12/6/6/6) and Grin1-KD (n = 8/6/8/6) for vehicle, haloperidol, UNC9994 and Haloperdiol + UNC9994 (Halo+UNC) groups, respectively. * p < 0.05; ** - p < 0.01; *** - p < 0.0001 in comparison with MK-801+vehicle-treated group or vehicle-treated Grin1-KD mice; ## - p < 0.01; ### - p < 0.001 – in comparison with vehicle-treated WT mice;

#### 3.1.2. Grin1-KD genetic model

The total travelled distance was significantly affected by genotype [F (1, 52) = 406.3; p < 0.0001], drugs [F (3, 52) = 11.5; p < 0.0001], genotype x drugs interactions [F (3, 52) = 9.2; p < 0.0001], time intervals [F (11, 572) = 96.4; p < 0.0001], and time intervals x genotype [F (11, 572) = 44.8; p < 0.0001]. Vehicle-treated Grin1-KD mice showed hyperactivity in comparison with vehicle-treated WT littermates in all tested time intervals (p’s < 0.001) (Figure 1C). UNC9994 and haloperidol modestly ameliorated hyperactivity of Grin1-KD mice (p’s < 0.05-0.01). Co-administration of UNC9994 with haloperidol substantially reduced their locomotor agitation in the open field (p’s < 0.001).

### 3.2. Effects of UNC9994 (0.25 mg/kg), haloperidol (0.15 mg/kg) and their co-administration on sensorimotor gaiting deficit in MK-801 (0.15 mg/kg) treated and in Grin1-KD mice

#### 3.2.1. MK-801 pharmacological model

The percentage of PPI was influenced by pre-pulses [F 2, 86) = 166.5; p < 0.0001], MK-801 [F (1, 43) = 50.5; p < 0.0001], drug treatment [F (3, 43) = 16.1; p < 0.0001], MK-801 x drugs interactions [F (3, 43) = 8.9; p < 0.0001], pre-pulses x MK-801 [F (2, 86) = 12.0; p < 0.0001], and pre-pulses x drugs [F (6, 86) = 4.4; p < 0.001] interactions. MK-801-treated mice expressed deficient PPI at all pre-pulses (p’s < 0.05-0.01) as compared with vehicle-treated animals (Figure 2A-B). Co-administration of haloperidol with UNC9994 efficiently corrected MK-801-induced PPI impairments (p’s < 0.01-0.001).

**Figure 2.**
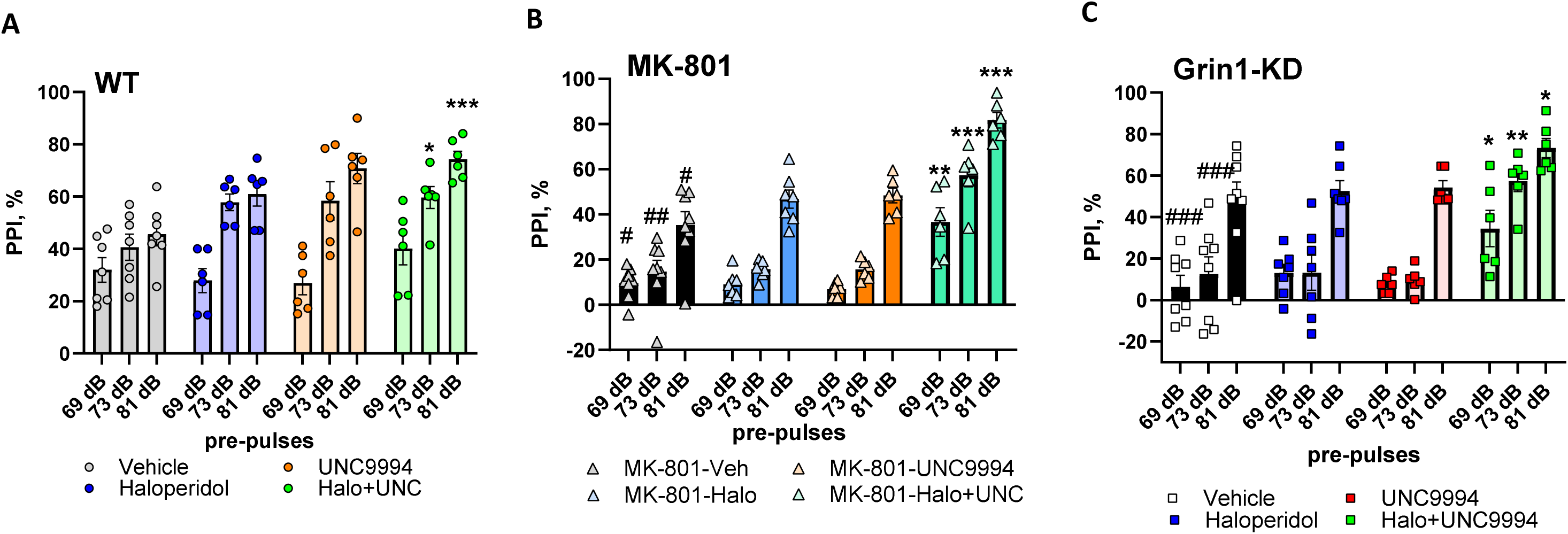
**A-C.** Effects of haloperidol (0.15 mg/kg), UNC9994 (0.25 mg/kg), and their co-administration on sensorimotor gaiting assessed as pre-pulse inhibition (PPI) of the acoustic startle response in vehicle-/WT (**A**), MK-801 (0.15 mg/kg)-treated wild-type (WT) mice (**B**), and Grin1 knockdown (Grin1-KD) mice (**C**). The percentage of PPI (PPI, %) is presented for each experimental group: WT (n = 7/6/6/6); MK-801-treated mice (n = 8/7/6/6) and Grin1-KD (n = 8/7/6/6) for vehicle, haloperidol, UNC9994 and Haloperdiol + UNC9994 (Halo+UNC) groups, respectively. * p < 0.05; ** - p < 0.01; *** - p < 0.0001 in comparison with MK-801+vehicle-treated group or vehicle-treated Grin1-KD mice; ## - p < 0.01; ### - p < 0.001 – in comparison with vehicle-treated WT mice;

There was a main effect of MK-801 treatment [F (1, 43) = 56.3; p < 0.05] but no effects of drug treatments or their interactions on the acoustic startle response (ASR) intensity (all p’s > 0.05) (Table 1).

**Table 1.**
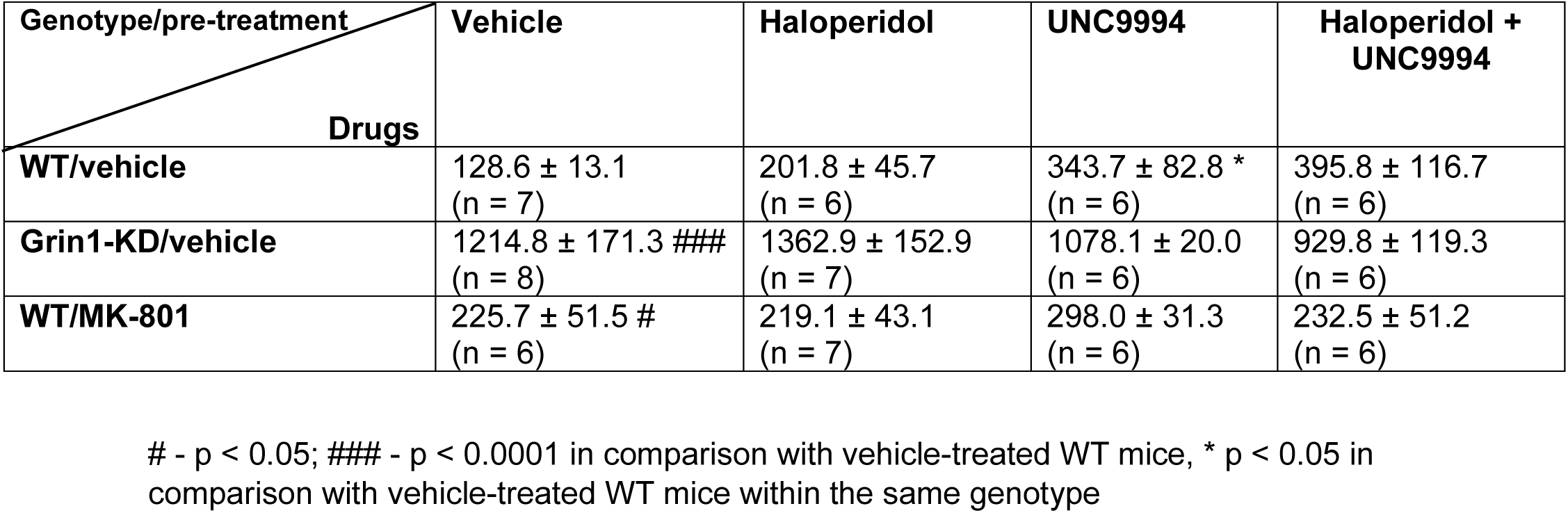
Effects of UNC9994 (0.25 mg/kg) and haloperidol (0.15 mg/kg) on the Acoustic Startle Response.

#### 3.2.2. Grin1-KD genetic model

PPI was substantially affected by pre-pulses [F (2, 86) = 95.0; p < 0.0001], genotype [F (1, 43) = 29.9; p < 0.0001], drugs [F (3, 43) = 6.4; p < 0.001], genotype x drugs [F (3, 43) = 5.5; p < 0.01], pre-pulses x genotype [F (2, 86) = 12.9; p < 0.0001], pre-pulses x drugs [F (6, 86) = 2.7; p < 0.05] and pre-pulses x genotype x drugs [F (6, 86) = 2.4; p < 0.05] interactions. The post-hoc analysis found that haloperidol co-administered with UNC9994 significantly facilitated PPI at all three pre-pulses in Grin1-KD mice (p’s < 0.05-0.01) and at 73 dB and 81 dB in WT animals (p < 0.05 and p < 0.001, respectively) (Figure 2A-C).

Measuring the startle response, two-way ANOVAs detected a main effect of genotype [F (1, 43) = 115.2; p < 0.0001] and genotype x drug interactions [F (3, 43) = 3.2; p < 0.05]. Vehicle-treated Grin1-KD mice showed the enhanced startle (p < 0.001) in comparison to WT mice (Table 1). UNC9994 modestly increased ASR in WT mice (p < 0.05) but there was no effect of haloperidol, UNC9994 or their combination on the ASR in any other experimental groups (p > 0.05).

### 3.3. Effects of UNC9994 (0.25 mg/kg), haloperidol (0.15 mg/kg) and their co-administration on the Spontaneous Alterations deficit and Repetitive Behavior assessed in Y-maze in MK-801 (0.15 mg/kg) treated and in Grin1-KD mice

#### 3.3.1. MK-801 pharmacological model

Assessing the percentage of spontaneous alterations (SA,%), two-way ANOVA detected a main effect of MK-801 treatment [F (1, 47) = 45.8; p < 0.001], drug treatments [F (3, 47) = 19.1; p < 0.0001] and MK-801 x drug treatments [F (3, 47) = 7.0; p < 0.001]. The post-hoc analysis found that co-administration of haloperidol with UNC9994 significantly facilitated working memory in Y-maze (p < 0.001) (Figure 3B), whereas all compounds administered alone did not affect MK-801-treated and vehicle-treated animals (Figure 3A-B). In addition, there was no synergistic effect between haloperidol and UNC9994 on working memory in vehicle-treated mice.

**Figure 3.**
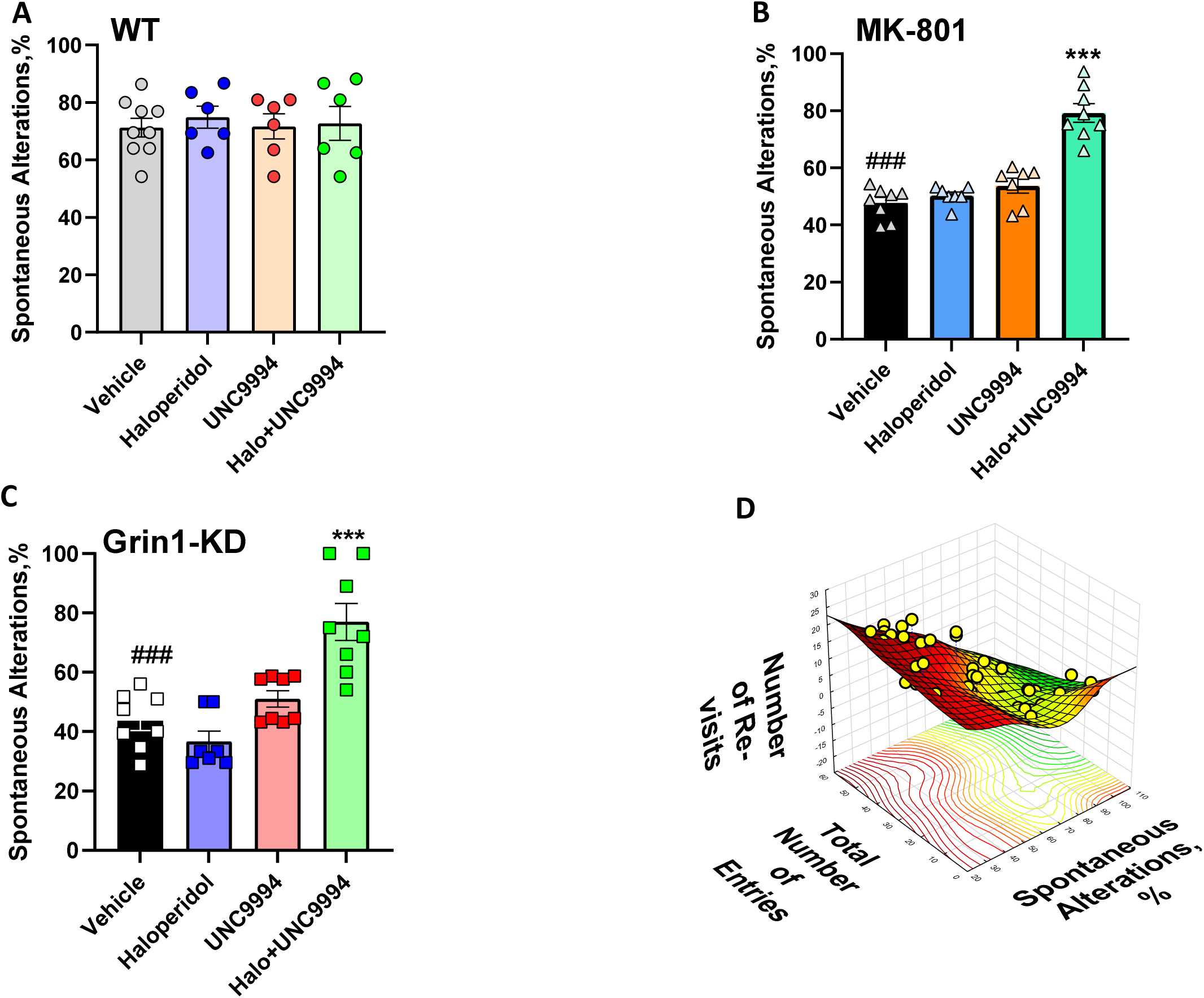
**A-D:** Effects of haloperidol (0.15 mg/kg), UNC9994 (0.25 mg/kg), and their co-administration on the percentage of spontaneous alterations (%) assessed in Y-maze in vehicle-/WT (**A**), MK-801 (0.15 mg/kg)-treated wild-type (WT) mice (**B**), and Grin1 knockdown (Grin1-KD) mice (**C**). WT (n = 9/6/6/6); MK-801-treated mice (n = 8/7/7/8) and Grin1-KD (n = 8/7/8/8) for vehicle, haloperidol, UNC9994 and Haloperdiol + UNC9994 (Halo+UNC) groups, respectively. * p < 0.05; ** - p < 0.01; *** - p < 0.0001 in comparison with MK-801+vehicle-treated group or vehicle-treated Grin1-KD mice; ## - p < 0.01; ### - p < 0.001 – in comparison with vehicle-treated WT mice. **D.** The number of re-visits of the same arm negatively correlates with the SA,%, and positively linked to the total number of entries (all Means ± SEM are presented in Table 2). The 3D surface graph illustrates surfaces fitted by a smoothing technique to the SA, %, total number of entries and number of re-visits for Grin1-KD and WT experimental mice. The color spectrum (from green to brown) represents the meaning of studied parameters (from low to high, respectively). Pearson correlation coefficients are: r = -0.76; p < 0.001 – for SA,% and number of re-visits; r = 0.66; p < 0.001 – for the total number of entries and number of re-visits; and r = -0.65; p < 0.01 for the total number of entries and SA, %. (N = 58).

**Table 2.** Effects of UNC9994 (0.25 mg/kg) and haloperidol (0.15 mg/kg) on the number of re-visits and the total number of entries in Y-maze.

| Genotype/pre-treatment<br>Drugs | Vehicle | Haloperidol | UNC9994 | Haloperidol +<br>UNC9994 |
| --- | --- | --- | --- | --- |
| **Number of re-visits** |  |  |  |  |
| **WT/vehicle** | 1.6 ± 0.6<br>(n = 9) | 1.7 ± 0.7<br>(n = 6) | 1.3 ± 0.5<br>(n = 6) | 2.2 ± 1.1<br>(n = 6) |
| **Grin1-KD/vehicle** | 15.5 ± 1.5 ###<br>(n = 8) | 15.4 ± 1.2<br>(n = 7) | 15.1 ± 1.1<br>(n = 8) | 6.5 ± 0.9 ***<br>(n = 8) |
| **WT/MK-801** | 12.1 ± 1.4 ##<br>(n = 7) | 12.1 ± 1.6 *<br>(n = 6) | 12.7 ± 1.3<br>(n = 6) | 6.9 ± 0.8 ***<br>(n = 6) |
| **Total number of entries** |  |  |  |  |
| **WT/vehicle** | 22.9 ± 1.6<br>(n = 9) | 19.3 ± 1.3<br>(n = 6) | 21.5 ± 2.2<br>(n = 6) | 21.0 ± 2.0<br>(n = 6) |
| **Grin1-KD/vehicle** | 49.3 ± 2.2 ###<br>(n = 8) | 39.4 ± 2.7 *<br>(n = 7) | 41.5 ± 3.9<br>(n = 8) | 21.3 ± 4.3 ***<br>(n = 8) |
| **WT/MK-801** | 37.5 ± 1.5 ##<br>(n = 7) | 31.1 ± 3.1 *<br>(n = 6) | 30.3 ± 1.8<br>(n = 6) | 22.5 ± 2.7 **<br>(n = 6) |

Measuring the number of re-visits, two-way ANOVA found a main effect of MK-801 treatment [F (1, 47) = 148.9; p < 0.001], drugs treatment [F (3, 47) = 3.1; p < 0.05] and MK-801 x drugs interactions [F (3, 47) = 3.2; p < 0.05]. MK-801-treated mice expressed repetitive behaviour, re-visiting the same arm of the maze significantly more than vehicle-treated animals (p < 0.0001) (Table 2). Dual administration of haloperidol with UNC9994 was able to ameliorate this rigid behaviour and reduced the number of re-visits (p < 0.0001) in MK-801-treated animals. Haloperidol or UNC9994 given alone had no effect on MK-801 or vehicle-pre-treated animals animals.

Measuring the number of entries, two-way ANOVA detected a main effect of MK-801 [F (1, 47) = 23.1; p < 0.01], drug treatments [F (3, 47) = 7.4; p < 0.01] and their interactions [F (3, 47) = 5.7; p < 0.05]. MK-801-treated mice expressed hyperactivity (p < 0.01), and haloperidol given alone (p < 0.05) or in combination with UNC9994 (p < 0.01) ameliorated hyperactivity (Table 2).

Pearson’s analysis found a negative correlation between number of re-visits and SA,% (r = -0.62; p < 0.001), between the total number of entries and SA,% (r = -0.45; p < 0.01) and positive correlation was found between number of re-visits and total number of visits of Y-maze’ arms (r = 0.53; p < 0.01).

#### 3.3.2. Grin1-KD genetic model

Assessing the percentage of spontaneous alterations, two-way ANOVA detected a main effect of genotype [F (1,50) = 45.7; p < 0.0001], drugs [F (3,50) = 8.0; p < 0.001] and their interactions [F (3,50) = 8.5; p < 0.0001]. Vehicle-treated Grin1-KD mice expressed deficient spontaneous alterations compared to WT mice (p < 0.001), which was corrected by the combined administration of haloperidol with UNC9994 (p < 0.001) but not by haloperidol or UNC9994 given alone (Figure 3C).

The number of re-visits was significantly affected by genotype [F (1, 50) = 245.9; p < 0.001], drugs [F (3, 50) = 7.8; p < 0.001], and genotype x drugs [F (3, 50) = 10.4; p < 0.0001] interactions. Vehicle-treated Grin1-KD animals expressed profound repetitive behaviour, re-visiting the same arm of the maze more often than vehicle-treated WT mice (p < 0.0001) (Table 2). Co-administration of haloperidol with UNC9994 reduced the number of re-visits in Grin1-KD mice (p < 0.001). Studied compounds given alone had no effects on this parameter in Grin1-KD mice and their WT littermates.

Measuring the total number of entries, two-way ANOVA detected a main effect of genotype [F (1, 50) = 47.8; p < 0.001], drug treatment [F (3,50) = 11.5; p < 0.001] and their interactions [F (3, 50) = 8.1; p < 0.001]. Vehicle-treated Grin1-KD mice showed hyperactivity (p < 0.001 vs vehicle-treated WT mice), which was slightly reduced by haloperidol (p < 0.05) and a more significant effect was seen after dual co-administration of UNC9994 with haloperidol (p < 0.001) (Table 2).

There was a negative correlation between the percentage of spontaneous alterations (SA, %) and number of re-visits (r = -0.76; p < 0.001), as well as between the total number of entries and SA,% (r = -0.65; p < 0.001). A positive association was detected between the number of re-visits and the total number of visits to Y-maze’s arms (r = 0.66; p < 0.001).

### 3.4. Effects of UNC9994 (0.25 mg/kg), haloperidol (0.15 mg/kg) and their co-administration on the deficit of the executive function assessed in the Puzzle box in MK-801 (0.15 mg/kg) treated and in Grin1-KD mice

#### 3.4.1. MK-801 pharmacological model

Measuring the latency to reach a goal box, two-way ANOVAs with repeated measures detected a main effect of trials [F (5, 205) = 76.8; p < 0.0001], interactions between trials and MK-801 [F (5, 205) = 4.9; p < 0.001], trials x co-administrated drugs [F (15, 205) = 5.9; p < 0.0001], and trials x MK-801 x co-administrated drugs [F (15, 205) = 8.8; p < 0.0001].

MK-801-treated mice showed impaired ability to solve the bedding obstacle on trials T5 and T6 (p’s < 0.001) (Figure 4B) in comparison with vehicle-treated animals (Figure 4A). Co-administration of either haloperidol alone or UNC9994 did not affect MK-801-induced impairments on T5 and T6 (p’s > 0.05). Co-administration of haloperidol with UNC9994 markedly facilitated executive functions on T5 and T6 in MK-801-treated animals (p’s < 0.001; Figure 4B).

**Figure 4.**
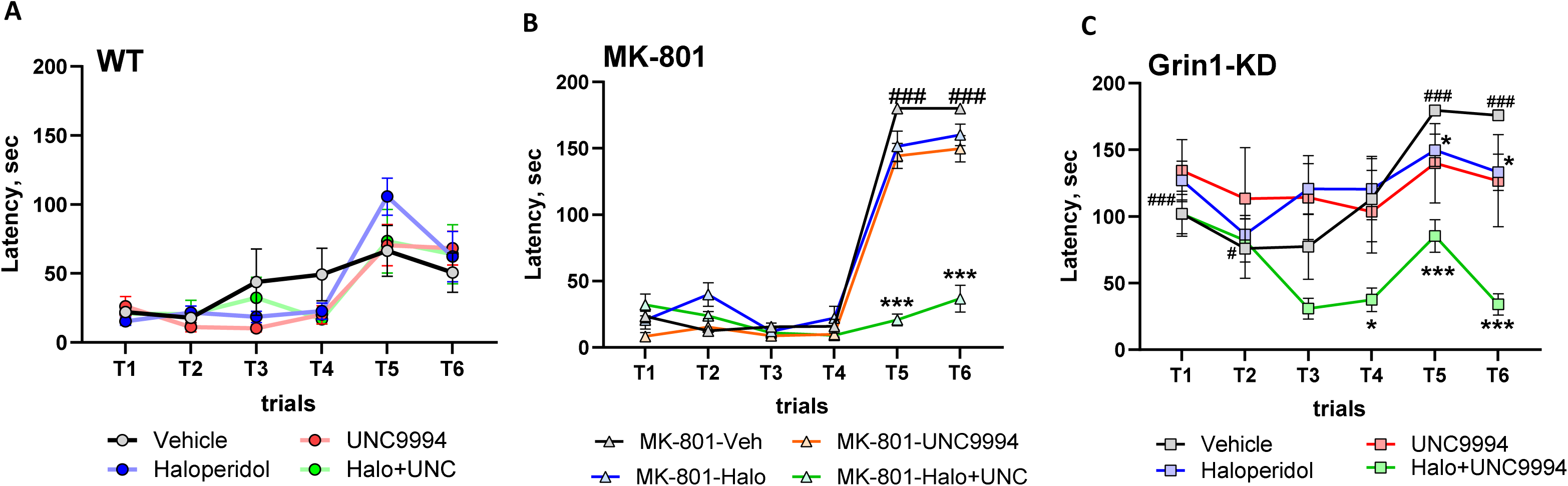
**A-C:** Effects of haloperidol (0.15 mg/kg), UNC9994 (0.25 mg/kg), and their co-administration on executive function, assessed in the Puzzle box paradigm. **A-C.** Latencies scored to reach the goal zone during the 6 trials of the test in vehicle-/WT (**A**), MK-801 (0.15 mg/kg)-treated wild-type (WT) mice (**B**), and Grin1 knockdown (Grin1-KD) mice (**C**). WT (n = 6/6/7/6); MK-801-treated mice (n = 6/6/6/6) and Grin1-KD (n = 6/6/6/6) for vehicle, haloperidol, UNC9994 and Haloperdiol + UNC9994 (Halo+UNC) groups, respectively. * p < 0.05; *** - p < 0.0001 in comparison with MK-801+vehicle-treated group or vehicle-treated Grin1-KD mice; ### - p < 0.001 – in comparison with vehicle-treated WT mice

#### 3.4.2. Grin1-KD genetic model

The latency to reach a goal box was significantly affected by trials [F (5, 200) = 23.9; p < 0.00001], genotype [F (1, 40) = 66.5; p < 0.0001], drugs [F (3, 40) = 3.4; p < 0.05], genotype x drug interactions [F (3, 40) = 3.2; p < 0.05], trials x genotype [F (5, 200) = 2.6; p < 0.05], trials x drugs [F (15, 200) = 2.1; p < 0.01], and trials x genotype x drugs [F (15, 200) = 3.3; p < 0.0001] interactions.

Vehicle-treated Grin1-KD mice expressed deficient performance on T1 (p < 0.001), T2 (p < 0.05) and solving the bedding obstacle on T5 and T6 (p’s < 0.001). Haloperidol given alone was able to modestly improve the performance of Grin1-KD mice on T5 and T6 (p’s < 0.05). Co-administration of haloperidol and UNC9994 significantly improved the performance of Grin1-KD mice in the Puzzle box, facilitating their long-term memory on T4 (p < 0.05), and executive ability to effectively solve bedding obstacles on T5 and T6 (p < 0.001) (Figure 4C). Haloperidol and UNC9994 administered alone or in combination did not affect the behavior of WT mice in the Puzzle box (Figure 4A).

## 4. Discussion

Our study revealed the strong efficacy of the combined administration of haloperidol and an analogue of aripiprazole, UNC9994, given in low doses, on schizophrenia-related behavioural phenotypes in pharmacological and genetic models of NMDAR hypofunction. The typical APD and UNC9994 given together, elicited beneficial effects on psychomotor agitation, and PPI deficit, phenotypes related to the positive symptoms of schizophrenia, as well as cognitive phenotypes in both animal models. Haloperidol + UNC9994 demonstrated efficacy on working memory and executive function in an NMDAR-dependent manner but also affected motor functions and sensorimotor gating in vehicle-pre-treated animals, regardless of MK-801 treatment or Grin1 genetic deficiency.

### Effects of UNC9994, haloperidol and their combination on Hyperactivity induced by NMDAR deficiency

Hyperactivity is a robust phenotype of Grin1-KD mice [36] and MK-801-treated animals (Mabunga et al., 2019; Jentsch et al., 1999; Gunduz-Bruce et al., 2009). Construct validity for hyperactivity induced by the NMDAR hypofunction stems from the effect of dissociative anaesthetics, such as ketamine and phencyclidine, to induce hallucinations with similarities to the positive symptoms of schizophrenia (Moghaddam et al., 2003; Tamminga et al., 2003). In rats and mice, moderate doses of ketamine and phencyclidine induce locomotor hyperactivity, but at higher doses, lead to ataxia and stereotyped behaviour, reducing locomotion (Mabunga et al., 2019; Wu et al., 2005). In our experiment low dose of MK-801 (0.15 mg/kg) induced hyperactivity in WT mice, gradually increasing the travelled distance after 15 minutes, reaching the maximum level between 30-40 minutes, and for the rest of the testing period, supporting previous studies (Mabunga et al., 2019). Genetic deficit of the Grin1 gene that reduces expression of the GluN1 subunit of NMDAR by ∼ 90% (Mohn et al., 1999), also leads to persistent hyperactivity (Mielnik et al., 2021; Milenkovic et al., 2014).

UNC9994 and haloperidol given alone in low doses had a modest effect on locomotor hyperactivity in both models, MK-801-treated animals, and Grin1-KD mice, supporting previous studies (Duncan et al., 2006; Park et al., 2016). Co-administration of the low dose UNC9994 with haloperidol reduced psychomotor to a larger degree in both models without showing any effect on WT animals. While Grin1-KD mice have been used to study APDs, including UNC9994 compound (Mohn et al., 1999; Duncan et al., 2006; Boulay et al., 2010; Park et al., 2016), our study is the first to assess the effects of the *polypharmacy* with UNC9994 and haloperidol on Grin1-KD mice and MK-801 pharmacological model. A few studies have reported that atypical but not typical APDs can reverse the behavioural effects of NMDAR antagonists which we also achieve with haloperidol with UNC9994 combination at low doses suggesting that co-administration of haloperidol and UNC9994 might elicit its effect via signalling pathways similar to atypical APDs.

### Effects of UNC9994, haloperidol and their combination on Cognitive phenotypes induced by NMDAR deficiency

Haloperidol and UNC9994 given alone at the studied doses did not affect cognitive behaviours, whereas their dual administration induced robust cognitive facilitation in PPI, working memory, and executive function.

#### Sensorimotor gating

PPI is a well-established endophenotype of schizophrenia with good face, predictive, and construct validity. Deficits in PPI are reported in schizophrenia (Swerdlow wt al., 1994; Braff et al., 2001) and in rodents, PPI is disrupted by NMDAR antagonists (Geyer et al., 2001), including low doses of MK-801 (Lipina et al., 2005). Atypical APDs are more efficacious than typical APDs for correcting the PPI deficit induced by NMDAR hypofunction (Yamada et al., 1999). We found that co-administration of haloperidol with UNC9994 at low doses significantly ameliorated PPI deficit at all three pre-pulses in both models without affecting the overall startle response, again suggesting that the co-administration of low dose haloperidol with UNC9994 might act similar to atypical-like APDs.

#### Working memory and executive functions

Human executive functions can be generally described as a complex of different high– level cognitive processes that enable individuals to regulate their thoughts and actions during a goal–directed behaviour by influencing lower–level processes (Friedman and Miyake, 2017). These functions include planning, working memory processes, switching between tasks or response inhibition, and their general role is to effectively implement goal–directed actions as well as to control attentional resources (Yogev-Seligmann et al., 2008). Schizophrenia is associated with deficits in executive functions, including impairments of working memory or behavioural flexibility (Wobrock et al., 2009).

#### Y-maze

In rodents, spontaneous alternation behaviour measured in Y-maze requires attention (Katz and Schmaltz, 1980) and working spatial memory. Normally, mice investigate a different arm of the maze rather than revisiting a previously explored one, indicative of sustained cognition. Although deficient spontaneous alterations have been already reported for Grin1-KD mice (Milenkovic et al., 2014; Chen et al., 2018) and MK-801-treated mice (Mabunga et al., 2019), a more detailed behavioural analysis in Y-maze was still missing in these two models. We found that both models showed severe cognitive rigidity, and more often re-visited the same arms in Y-maze than control animals. Moreover, there was a negative correlation between the percentage of spontaneous alterations and the number of re-visits, highlighting the repetitive behaviour as a major reason for the impaired performance of Grin1-KD and MK-801-treated mice in Y-maze. These findings are in good agreement with a recent study (Cleal et al., 2021), where repetitive visits of the previously visited arm were detected as a robust index of cognitive flexibility in Y-maze across a range of experimental conditions and multiple species, including zebrafish, mice, *Drosophila*, and humans. Our results show that a combination of haloperidol with UNC9994 at low doses was able to ameliorate the impaired behavioural performance in Y-maze in both studied models correcting the number of re-visits highlighting combination of haloperidol with UNC9994 as a new approach to correct rigid behaviour.

#### Puzzle Box

The Puzzle box was recommended as a relevant and faster way to assess executive functions in a mouse (Ben Abdallah et al., 2011) as compared to the attentional set-shifting (Garner et al., 2006) or the 5-choice serial reaction time (Robbins, 2002) which require long training. It has been shown that Grin1-KD mice have impairments in this paradigm (Mielnik et al., 2021; Milenkovic et al., 2014; Islam et al., 2017), where altered somatosensory functions play an important role in affecting Puzzle box performance in these animals (Lipina et al., 2022). Our current experiments confirmed the Puzzle box impairments in Grin1-KD animals and haloperidol and UNC9994 given alone were not able to correct this deficit. As with the other observations, co-administration of both compounds significantly ameliorated the ability to solve the bedding obstacle on Trial 5 and Trial 6 and facilitated a long-term memory on Trial 4. Importantly, we demonstrated, for the first time, that acute administration of MK-801 severely affected the ability of the experimental mice to solve the bedding problem on Trial 5 and Trial 6 of the Puzzle box, similar to Grin1-KD mice. Again, this impairment observed in MK-801 treated animals was reversed by the dual pharmacological treatment with haloperidol and UNC9994. The deficits we have observed in the Puzzle box paradigm in both of our models, Grin1-KD and MK-801 treated animals, denotes the importance of intact NMDAR signaling on executive function for this assay.

### Polypharmacy between UNC9994 and haloperidol: mechanisms and future

UNC9994 was discovered as a unique β-arrestin-biased functionally specific to the D2R compound (Allen et al., 2011). It acts as a partial agonist that stimulates D2R-mediated β--arrestin recruitment and signalling, and at the same, it is inactive at the Gi-dependent pathway. Despite UNC9994 being an agonist for β-arrestin-2 recruitment, in contrast to quinpirole it does not trigger robust D2R internalization (Allen et al., 2011). The reduced internalization is beneficial as it may prevent side effects such as tachyphylaxis and receptor down-regulation (Allen and Roth, 2011). Another unique feature of the UNC9994 agent is that it is more efficacious (Emax = 91 ± 3%) than aripiprazole (Emax = 73 ± 1%) to act as a partial agonist for β-arrestin-2 recruitment to D2R (Allen et al., 2011). Haloperidol does not activate D2R-mediated β-arrestin-2 translocation (Granger and Albu, 2005). The addition of haloperidol to UNC9994 may strongly inhibit D2R and subsequently reduce locomotor hyperactivity. From a pharmacological perspective, when UNC9994 is added to haloperidol, three substances compete to bind dopamine D2R: UNC9994 (a partial agonist of D2R (Ki = 0.75 nM; with slow dissociation koff = 31 min), haloperidol (Ki = 0.57 nM; koff = 1 min), and dopamine (Ki = 16 nM; koff = 24 min) (Allen et al., 2011; Pigott et al., 2003; Sánchez-Soto et al., 2016; de Witte et al., 2018). Hence, when UNC9994 is added to haloperidol, it could replace haloperidol from D2R sites and exert its partial agonist intrinsic activity. As a result, co-administration of UNC9994 with haloperidol could ameliorate negative symptoms and facilitate cognitive deficits as detected in our pre-clinical NMDAR hypofunctional models. Indeed, it has been shown that compounds with partial D2R agonistic activity induced cognitive enhancement in various pre-clinical studies (Kolaczkowski et al., 2015; Thompson et al., 2016).

Importantly, the balanced functionality of the NMDAR is principally required in the cortex to regulate executive functions and cognitive flexibility. The complex interactions between glutamatergic and dopaminergic systems critically modulate the gating of information flow in a brain and their impaired interactions lead to several psychopathologies, including schizophrenia (West et al., 2003). Pharmacological inhibition of NMDAR by non-competitive antagonist MK-801 affects not only glutamatergic transmission but also dopaminergic release (Bartsch et al., 2015; Saoud et al., 2021; Tsukada et al., 2005), indirectly activating the dopaminergic functions in humans (Kegeles et al., 2000) and rats (Adams et al., 1998). Genetic knockdown of the Grin1 gene significantly affects the dopaminergic functions (Ferris et al., 2014), leading to desensitization of the D2R, a faster spontaneous firing rate of the dopaminergic neurons, accompanied by the attenuated dopamine synthesis and release with increased dopamine clearance. A recent study by Thompson et al. (2016) showed that high dopamine concentration or D1R-D2R agonist can target cortical dopamine receptors coupled with a phospholipase C (PLC) to enhance excitatory synaptic transmission and improve behavioural performance of juvenile rates in the reversal-learning task (Thompson et al., 2016). Hence, the combination of UNC9994 with haloperidol may elicit their beneficial effects on cognitive flexibility of the NMDAR deficient mice in our study through activation of cortical D1R-D2R complex, which can be further explored in the future.

## 5. Conclusions

Taken together, our pre-clinical experiments show that dual administration of haloperidol and UNC9994 in low doses ameliorates behavioural phenotypes related to positive, negative, and cognitive symptoms of schizophrenia, supporting polypharmacy as a new effective approach for the treatment of schizophrenia and development of new APDs. Further validation of the efficacy of the combined treatment to prevent schizophrenia-related behavioural phenotypes in additional genetic mouse models is needed to generate optimal strategies for translational studies in humans.

## Author Contributions

Conceptualization, funding acquisition, validation, original draft editing, project administration, supervision: A.J.R, A.S., W.W. Methodology, performance of experiments, data acquisition and analysis, original draft preparation: T.L. Providing UNC9994 compound: W.W. All authors have approved the submitted manuscript prior to submission. All authors agree to be personally accountable for their own contributions and for ensuring that questions related to the accuracy or integrity of any part of the work, even ones in which they were not personally involved, are appropriately investigated, resolved, and documented in the literature. All authors have read and agreed to the published version of the manuscript.

## Informed Consent Statement

Not applicable.

## Data Availability Statement

Detailed statistical analysis and raw data can be obtained upon request to the authors.

## Acknowledgements

We thank Dr. Jian Jin at the University of North Carolina in Chapel Hill for providing UNC9994 compound and Wendy Horsfall for genotyping of the experimental mice.

